# A removable and cosmopolitan dsRNA Toti-like virus causes latent infection in a model diatom strain

**DOI:** 10.1101/2024.01.12.575345

**Authors:** Jiahuan Zhang, Chenjie Li, Xiaofeng Xiong, Kangning Guo, Yanlei Feng, Huan Zhang, Hanhua Hu, Xiaobo Li

**Affiliations:** College of Life Sciences, Zhejiang University, Hangzhou, Zhejiang, China; School of Life Sciences, Westlake University, Hangzhou, Zhejiang, China; Key Laboratory of Algal Biology, Institute of Hydrobiology, Chinese Academy of Sciences, Wuhan, China; University of Chinese Academy of Sciences, Beijing, China; Hangzhou Global Scientific and Technological Innovation Center, Zhejiang University, Hangzhou, China; Institute of Biology, Westlake Institute for Advanced Study, Hangzhou, China

**Keywords:** Diatom, virus, *Phaeodactylum tricornutum*, *Totiviridae*, coat protein

## Abstract

Diatoms contribute to 20% of global primary productivity. Although some diatom viruses have been identified, the molecular mechanisms underlying their interactions with the host remain poorly understood. In this study, we report the discovery of an RNA molecule in the DNA extracts of the *Phaeodactylum tricornutum* strain Pt1, which possesses a well-annotated genome and has been used as a diatom model system since 1956. We confirmed this molecule to be a double-stranded linear RNA molecule and, through sequencing, demonstrated it to be a virus in the *Totiviridae* family that is prevalent among marine stramenopiles. We also detected this virus in *Phaeodactylum tricornutum* strain Pt3, which was collected in 1930s from a similar geographic location to Pt1, suggesting its prevalence within the region. By employing various inhibitors of the viral RNA-dependent RNA polymerase, we successfully generated a virus-free line isogenic to Pt1, establishing a model system to investigate the impact of RNA viruses on diatom physiology. The virus-free lines did not display obvious growth advantages or defects, indicating a tendency of the virus towards latent infection. Furthermore, we generated a robust antibody against the coat protein of this virus. By performing immunoprecipitation coupled with mass spectrometry, we found that translation-related proteins are enriched as potential interacting partners of the coat protein. Our results suggest that potential viral impacts in molecular research should be considered when Pt1 and Pt3 are used for studying translation-related processes. Additionally, our study unveiled a temperate mode of interaction between viruses and marine algal hosts that differs from the commonly-reported virulent, lytic infections.

**Highlights:**

1. prevalent dsRNA virus belonging to the *Totiviridae* family was discovered in the Pt1 and Pt3 strains of the model diatom *Phaeodactylum tricornutum*.
2. virus is absent in eight other strains of *P. tricornutum*, highlighting the importance of studying multiple accessions.
3. virus can be eliminated using a nucleotide analogue, resulting in a virus-free isogenic strain that allows us to investigate how viruses may affect diatom physiology.
4. robust antibody against the coat protein of this virus was developed to assist mechanistic studies of diatom-virus interactions.

## Introduction

Diatoms encompass more than one hundred thousand species [1]. They constitute a major group of unicellular photosynthetic eukaryotes. Additionally, they contribute approximately 20% of global photosynthetic carbon dioxide fixation and are thus important primary producers [2]. In the meanwhile, diatoms and related biogeochemical processes are also influenced by various organisms, such as viruses, bacteria, or animals [3].

In marine environments, viruses have been identified as the most abundant microorganisms (by number) [4]. Moreover, RNA viruses deeply and significantly influence marine ecology based on their disturbance of the metabolism of their phytoplankton hosts [4]. The hosts of RNA viruses in the marine system, especially microalgae, hold irreplaceable value in aquaculture, pharmaceutical industry, and bioenergy [5–7]. Virus infection in microalgae could impair biofuel production during industrial cultivation [8]. Although the impact of virus infection on marine microalgae hosts is considered profound and comprehensive, progress in this research field has been slow. Despite the development and improvement of metagenomics, which has greatly facilitated the discovery of new viruses in marine microalgae [9–11]. However, the hosts of the vast majority of the viruses are either unknown, or not culturable in the lab. These challenges have hampered in-depth research on the effects of virus infection and the interaction between viruses and microalgae hosts. The establishment of a genetically tractable model system for studying diatom viruses and their relationship with their hosts remains a gap in this research field.

In previous studies on viral infections in diatoms, only a few single-stranded DNA or RNA viruses have been relatively well-studied [12, 13]. These viruses were found to be lytic to the host diatoms, and the infections were not only confined to particular species but also specific to certain strains of the host [13]. However, research on double-stranded DNA or RNA viruses in diatoms has been limited. Moreover, the hosts used in these studies were not model strains of diatoms with well-sequenced genomes, which restricted further research on the interactions and influences between diatoms and viruses, especially double-stranded DNA or RNA viruses.

*Phaeodactylum tricornutum* (*P. tricornutum*) is a prevalent coastal diatom. It is a promising source of value-added products like eicosapentaenoic acid and fucoxanthin [14]. It has been used as a model system since 1910 when a strain was collected near Blackpool in UK [15]. It has a well-developed toolbox such as the chromosomal-level genome sequence (for Pt1) [16] and multiple methods (biolistic, electroporation, conjugation) for transformation [17–19]; this toolset is much advanced compared with other diatoms. Many strains of *P. tricornutum* have been collected and used for ecological and evolutionary research [20, 21]. In addition, it has been used for studies of various processes with biogeochemical implications such as photosynthesis [22–26], cell morphology control [21] and nutrient deprivation responses [27, 28]. It is worth mentioning that most of the scientific discoveries in *P. tricornutum* were based studies using Pt1. Here, we discovered a double-stranded RNA (dsRNA) Toti-like virus in the Pt1 strain of *P. tricornutum* [16] and investigated the occurrence of this virus across different accessions of this diatom species. Characterization of this diatom-virus interaction model revealed a latent mode of infection and caveats in current research in *P. tricornutum* carried out using a single strain.

## Materials and Methods

### Algal strains and culture conditions

Strains Pt1 (CCMP2561), Pt5 (CCMP630), Pt6 (CCMP631), Pt7 (CCMP1327) and Pt9 (CCMP633) originated from the Provasoli-Guillard National Center for Culture of Marine Phytoplankton (CCMP). Strains Pt2 (CCAP 1052/1A), Pt3 (CCAP 1052/1B) and Pt4 (CCAP 1052/6) originated from the Culture Collection of Algae and Protozoa (CCAP). Strain Pt8 (NEPCC 640) originated from the Canadian Center for the Culture of Microorganisms (CCCM) while strain Pt10 (MACC B228) originated from the Microalgae Culture Collection of Qingdao University (MACC). All *P. tricornutum* strains Pt1-Pt10 in Hanhua Hu Lab were gifts from Chris Bowler. The Pt1 strain in Xiaobo Li Lab was ordered from CCMP.

The algal cultures were grown in f/2 media under white fluorescent lights (60 µmol m^−2^ s^−1^) at a temperature of 20°C. Solid f/2 media contained 1% agar. A 12-hour light-dark cycle was maintained. Nucleic acid and protein extraction were performed when the cell density reached approximately 1 × 10^7^ cells/mL, as estimated using the Countess II cell counter (AMQAX1000, Thermofisher).

### Total nucleic acid extraction

For each sample, 5 × 10^8^ cells were harvested and resuspended in 1 mL cetyltrimethylammonium bromide (CTAB) extraction buffer (100 mM Tris, 1.4 M NaCl, 20 mM EDTA, pH 8.0, 2% CTAB, 0.3% β-mercantoethanol) [29]. After the incubation at 65°C for 1 hour, an equal volume of DNA extraction reagent (T0250, Solarbio) was used for extraction according to the instructions of the reagent. Then the aqueous phase was extracted by DNA/RNA Extraction Reagent (Chloroform:Isoamylol=24:1; P1014, Solarbio) twice according to the instructions of the reagent. The nucleic acids were precipitated by adding an equal volume of isopropanol and incubated at 4°C for 30 minutes. After centrifugation at 13,000 × g at 4°C, the supernatant was carefully removed without disturbing the pellet. The pellet was washed twice with 75% ethanol and dried. Finally, 50 µL of nuclease-free water was added to dissolve the pellet. The extracts from each sample were resolved by agarose gel electrophoresis.

### RNA extraction and purification

To perform nuclease S1 digestion of total nucleic acid samples, next-generation sequencing of dsRNA, and rapid amplification of cDNA ends, we harvested and dried out 2 × 10^9^ cells using vacuum filtration. The dried cells were then ground in liquid nitrogen using mortars. For total RNA extraction, 10% of the resulting powder was mixed with 1 mL of Trizol (15596026, Invitrogen). The total RNA extraction was carried out as previously reported [30, 31].

For nuclease S1 digestion, the RNA sample was treated with nuclease S1 (EN0321, Thermofisher) according to the product instructions and resolved by agarose gel electrophoresis.

For strand-specific next-generation sequencing of dsRNA, the RNA sample was digested using nuclease S1 (EN0321, Thermofisher) and DNase I (EN0521, Thermofisher) [32] according to the manufacturers’ instructions. The digested RNA was then purified using VATHS DNA Clean Beads (N411, Vazyme Biotech) to remove short digested fragments of nucleic acid.

For rapid amplification of cDNA ends (RACE), the dsRNA was purified from the total RNA sample using the Plant Viral dsRNA Enrichment Kit (5345, MBL Beijing Biotech) according to the manual.

### Next-generation sequencing of dsRNA

For strand-specific next-generation sequencing of dsRNA, the purified dsRNA sample was denatured at 99°C for 10 minutes, followed by rapid chilling on ice [33]. Subsequently, a library for strand-specific RNA sequencing (RNA-seq) was constructed using the VAHTS® Stranded mRNA-seq Library Prep Kit for Illumina V2 (NR612, Vazyme Biotech) with beads-purified dsRNA. The library was then sequenced by Novogene Inc. The obtained reads were subjected to trimming and filtering using the FASTp software [34]. The resulting clean reads from strand-specific cDNA sequencing were assembled using the Trinity software [35]. To identify the dsRNA sequence, the assembled sequences were selected if both complementary sequences were detected. The Integrative Genomics Viewer (IGV) software [36] was used for the visualization of the mapped reads.

### Determination of the terminal sequence of dsRNA by RACE

The purified dsRNA was used for RACE after the denaturation as described above. The RACE steps (**Fig. S1A**) were performed immediately following the denaturation of dsRNA, and the three most common sequences detected by next-generation sequencing from each terminal were presented in **Supplementary Table S2**. A DNA adaptor (P0) for RACE, with a phosphorylation (P0, **Supplementary Table S1**) at the 5’ terminal and a dideoxycytosine nucleotide (ddC) at the 3’ terminal, was synthesized in advance.

The sequences of the 3’ terminal ends of both the positive and negative strands were determined using RNA ligase-mediated rapid amplification of complementary DNA (cDNA) ends (RLM-RACE) [37]. Briefly, the denatured dsRNA was mixed with the DNA adaptor for adaptor ligation using T4 RNA ligase 1 (M0204, NEB). Subsequently, the adaptor-complementary primer P1 (**Supplementary Table S1**) was used for reverse transcription of the terminal of each strand using SuperScript™ III Reverse Transcriptase (18080093, Invitrogen) at 50°C as described by the product instruction. The resulting product was purified using VATHS DNA Clean Beads (N411, Vazyme Biotech) as described by the product instruction. Finally, the reverse transcription product was used in polymerase chain reaction (PCR) to construct a library for next-generation sequencing, using first PCR primers (P1, P2, or P3) and library construct PCR primers (P1, P4, or P5, **Supplementary Table S1**).

The 5’ ends of both the positive and negative strands were determined by using adapter-ligated RACE [38] with minor modifications. Firstly, the cDNA of the terminal of each strand was synthesized using SuperScript™ III Reverse Transcriptase (18080093, Invitrogen) and gene-specific primers (GSPs) P2 and P3 (**Supplementary Table S1**), followed by RNA digestion using RNase A (EN0531, Thermofisher) as described by the product instructions. Subsequently, the sample was purified using VATHS DNA Clean Beads (N411, Vazyme Biotech), and the DNA adaptor was added for cDNA adaptor ligation using CircLigase™ II ssDNA Ligase (CL9021K, Lucigen) as described by the product instructions. his enzyme catalyzes the ligation of single-stranded DNA (ssDNA) and single-stranded RNA (ssRNA) substrates that have a 5’-monophosphate and a 3’-hydroxyl group, respectively. The ligation product was then purified using VATHS DNA Clean Beads (N411, Vazyme Biotech) as described by the product instruction. Finally, the reverse transcription product was used in PCR to construct a library for next-generation sequencing, using first PCR primers (P1, P2, or P3) and library construct PCR primers (P1, P4, or P5, **Supplementary Table S1**).

### Reverse transcription-PCR and Sanger sequencing

For the verification of the coding region sequence on the dsRNA in Pt3, reverse transcription-PCR (RT-PCR) was performed. Briefly, the purified dsRNA from Pt3 was denatured as described above, followed by reverse transcription using SuperScript™ III Reverse Transcriptase (18080093, Invitrogen) and GSPs (P6 – P11, **Supplementary Table S1**) as described by the product instruction. The resulting product was then used as the template for PCR, using the same primers. Subsequently, the PCR product was subjected to Sanger sequencing using GSPs (P6 - P8, P10, P12-14, **Supplementary Table S1**).

### Removal of dsRNA using nucleotide analogues

The strategy of dsRNA depletion by nucleotide analogues was mainly based on the previous report on the elimination of dsRNA *Leishmania* RNA virus 1 (LRV1) in *Leishmania guyanensis* using nucleotide analogues [39]. In brief, 2’-C-Methyladenosine (2CMA; HY-125371, MedChemExpress) or Remdesivir (SF1193, Beyotime) were added into cells to a final concentration of 200 μM. The wild-type Pt1 in stationary phase (about 2 × 10^7^ cells/mL) was inoculated in new f/2 media with the dilution by 1000 times. Afterwards, the culture was grown to stationary phase. The 1000-fold dilution and the new round of culture were repeated at least five times. Then the culture was diluted and spread onto a f/2 agar plate (1% w/v). Finally, the clones were inoculated in new f/2 liquid media to verify the depletion of the dsRNA and coat protein (CP). The virus-free clones were named DR-strains while the virus-containing Pt1 clones were named DR+ strains.

### Protein extraction

Approximately 2 × 10^9^ algal cells were disrupted by grinding in liquid nitrogen, following the RNA extraction method described above, to extract intracellular proteins. For non-denaturing protein extraction, 5 mL of 1 × IP buffer [40] was added to the disrupted sample. After incubating at 4°C for 15 minutes, the cell debris were removed by centrifugation at 16,000 × g for 10 minutes. The supernatant was directly used or stored at −80°C.

For denatured protein extraction, the 1 × IP buffer was replaced with Laemmli Sample Buffer (1610747, Bio-Rad), and 1% 2-Mercaptoethanol was added to the sample. The sample was then boiled at 99°C for 5 minutes.

### Anti-CP antibody production

The production of anti-CP antibodies was carried out by ABclonal Technology Co., Ltd. (Wuhan). The full-length CP gene was cloned into the pET28a vector with a C-terminal 6×His-tag and expressed in *Escherichia coli* Rosetta (DE3) for subsequent immunization of rabbits.

The CP antigen was purified using a 5 mL HisTrap column (17524801, GE Healthcare) followed by a 5 mL HiTrap Q HP column (17115401, GE Healthcare) on AKTA Pure 25 system. Polyclonal antibodies against CP were obtained from rabbits immunized with the purified CP antigen.

The obtained antibodies were further purified using affinity chromatography. Purification was performed using columns containing purified full-length CP, while antibodies with non-specific affinity to host proteins were depleted using columns containing total proteins from the dsRNA-negative strain Pt1.

### Immunoprecipitation

Immunoprecipitation was performed following a previously described protocol with minor adjustments [40]. Approximately 1 mg of non-denatured total protein was mixed with 5 μg of CP antibody and incubated overnight at 4°C. After incubation, the protein A/G magnetic beads (88803, Pierce) were added and the mixture was incubated at room temperature for 2 hours. The samples were then treated according to the instruction of the beads and the protein complexes were then eluted by incubating at 99°C for 5 minutes with Laemmli Sample Buffer (1610747, Bio-Rad).

For the immunoprecipitation experiments, three virus-containing Pt1 clones were used, while three virus-depleted Pt1 clones were used as negative controls. Approximately 30% of each sample was analyzed by SDS-PAGE followed by staining with Coomassie Brilliant Blue G-250 (CBB).

### Immunoblot analysis

For each sample, 30-50 μg of total protein was boiled at 99°C for 5 minutes with the addition of Laemmli Sample Buffer (1610747, Bio-Rad) containing 1% 2-Mercaptoethanol in a total volume of 60-80 μl. The protein samples were separated on a gel prepared using the CFAS any KD PAGE protein electrophoresis gel preparation kit II (PE008-2, ZhongHuiHeCai) and transferred to a polyvinylidene difluoride (PVDF) membrane following the manufacturer’s protocol.

The membrane was blocked for 1 hour at room temperature with 5% skim milk in TBS buffer (80g/L NaCl, 2g/L KCl, 30g/L Tris base, pH 7.4) containing 0.05% Tween20. Subsequently, the membrane was incubated overnight at 4°C with the anti-coat-protein antibody (diluted 1:5000).

Afterwards, the membrane was washed three times with TBS-T for 10 minutes each, followed by incubation with the Goat anti-Rabbit IgG (H+L) HRP Conjugate Secondary Antibody (diluted 1:10000; HS101-01, TransGen Biotech) at room temperature for 1 hour. Before visualizing the protein bands, the membrane was washed five times with TBS-T for 10 minutes each.

### LC-MS/MS for co-immunoprecipitation

The denatured protein samples from co-immunoprecipitation were run into stacking gel and the sample gel band stained by Coomassie Brilliant Blue G250 were digested by trypsin (V5280, Promega) with prior reduction and alkylation in 50 mM ammonium bicarbonate at 37°C overnight. The digested products were extracted twice with 1% formic acid in 50% acetonitrile aqueous solution and dried to reduce volume using centrifugal evaporation.

The peptides were separated by a 135-min gradient elution at a flow rate of 0.3 µL/min using a Thermo Vanquish Neo integrated nano-HPLC system directly interfaced with a Thermo Exploris 480 mass spectrometer equipped with FAIMS Pro. The analytical column was a home-made fused silica capillary column (75 µm ID, 150 mm length; Upchurch) packed with C-18 resin (300 Å, 2 µm, Varian). Mobile phase A consisted of 0.1% formic acid in water, and mobile phase B consisted of 80% acetonitrile and 0.1% formic acid. The mass spectrometer was operated in data-dependent acquisition mode using Xcalibur 4.1 software, and the −45 V and −65 V CV values of FAIMS Pro were set as the two independent acquisition events. Under one event, a single full-scan mass spectrum in the Orbitrap (350-1800 m/z, 60,000 resolution) was followed by several data-dependent MS/MS scans (110-1500 m/z, 15,000 resolution) at 30% normalized collision energy, with a total cycle time of 1 s per event. The AGC target was set as 5e4, and the maximum injection time was 50 ms.

To conduct an all-round proteomic analysis, the clustered proteome reference database was generated using software CD-HIT [41] with parameter settings of −c 0.80 and −n 5 to cluster various reference databases (ASM15095v2 from NCBI and Ensembl, UP000000759 from Uniprot, NC_016739.1 from NCBI and two protein ORFs encoded by dsRNA virus genome). Each mass spectrum was analyzed using the Thermo Xcalibur Qual Browser and Proteome Discoverer (2.5.0.400) for the database searching against the clustered version of proteome database. The Sequest search parameters included a 10 ppm precursor mass tolerance, 0.02 Da fragment ion tolerance, and up to 2 internal cleavage sites. Fixed modifications included cysteine alkylation and the methionine oxidation was variable modification. Peptides were filtered with 1% false discovery rate (FDR).

Afterwards we selected out proteins with high confidence in peptide comparison (Protein FDR Confidence = High) and LC-MS/MS experiment (Exp. *Q*-value < 0.001) (**Supplementary Table S3**). The threshold of fold change between samples and controls was set to 8 while the threshold of *P*-value was set to 0.01 (**Supplementary Table S4**).

### Gene ontology (GO) analysis

The candidate list of co-immunoprecipitation was used for GO analysis. GO terms of *P. tricornutum* nuclear-encoded genes were predicted by InterProScan v5.28-67.0 [42] and Pannzer website [43]. GO enrichment analysis was performed by clusterProfiler v4 [44] in R-studio software.

## Results

### The presence of dsRNA Toti-like virus in Pt1

When analyzing nucleic acid extracts from the *P. tricornutum* strain Pt1, we observed an unexpected band at the position of the 5 kb DNA marker band on an agarose gel (**Fig. 1A**). It was evident that this nucleic acid was not genomic DNA, nor plastidic and mitochondrial DNA, as these molecules are all larger than 8 kb. We could also observe this band in the total RNA extract of *P. tricornutum* analyzed through agarose gel electrophoresis (**Fig. 1A**). To determine the identity of this band, we treated the total nucleic acid extract with DNase I and a separate total RNA extract with nuclease S1; the latter specifically degrades single-stranded DNA and RNA. The band at the 5 kb marker band position remained intact after either treatment (**Fig. 1A**), suggesting that this band is likely a double-stranded RNA (dsRNA) molecule. Furthermore, this nucleic acid was not only present in Pt1 cultured in Xiaobo Li’s lab (CCMP2561) but also in the strain of Pt1 in Hanhua HU’s lab (CCMP2561; **Fig. 1A and Fig. 2A**), suggesting that this nucleic acid is stably present in the Pt1 strain and is not a result of infection in one of our labs.

**Fig. 1.**
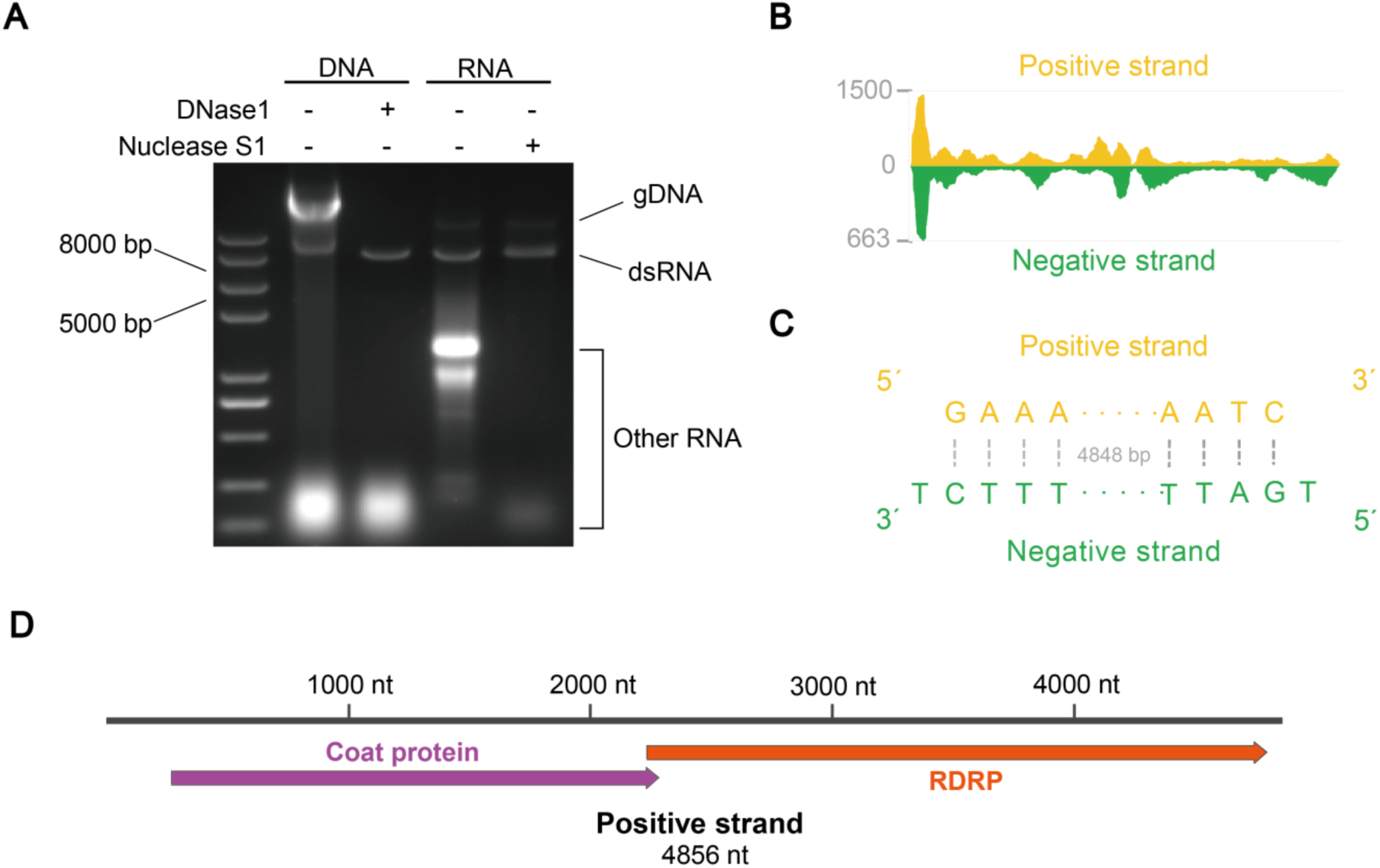
The discovery and determination of the complete genome of the dsRNA virus in the model diatom strain *P. tricornut*um CCMP2561. (A) Agarose gel image displaying the electrophoretic profile of DNA and RNA extracts treated with DNase I or Nuclease S1, resolved on a 1% agarose gel. (B) Integrative Genomics Viewer (IGV) snapshot depicting the coverage of the reads from strand-specific RNA-seq on both positive and negative strands of the dsRNA genome. The Y-axis represents the coverage of mapped reads. (C) The most abundant version of the terminal sequences of the dsRNA genome determined by RACE. (D) The scheme showing the protein-coding structure of the dsRNA positive strand. The Pt1 strain used in this panel is CCMP2561 maintained in Xiaobo Li Lab (see **Materials and Methods**).

**Fig. 2.**
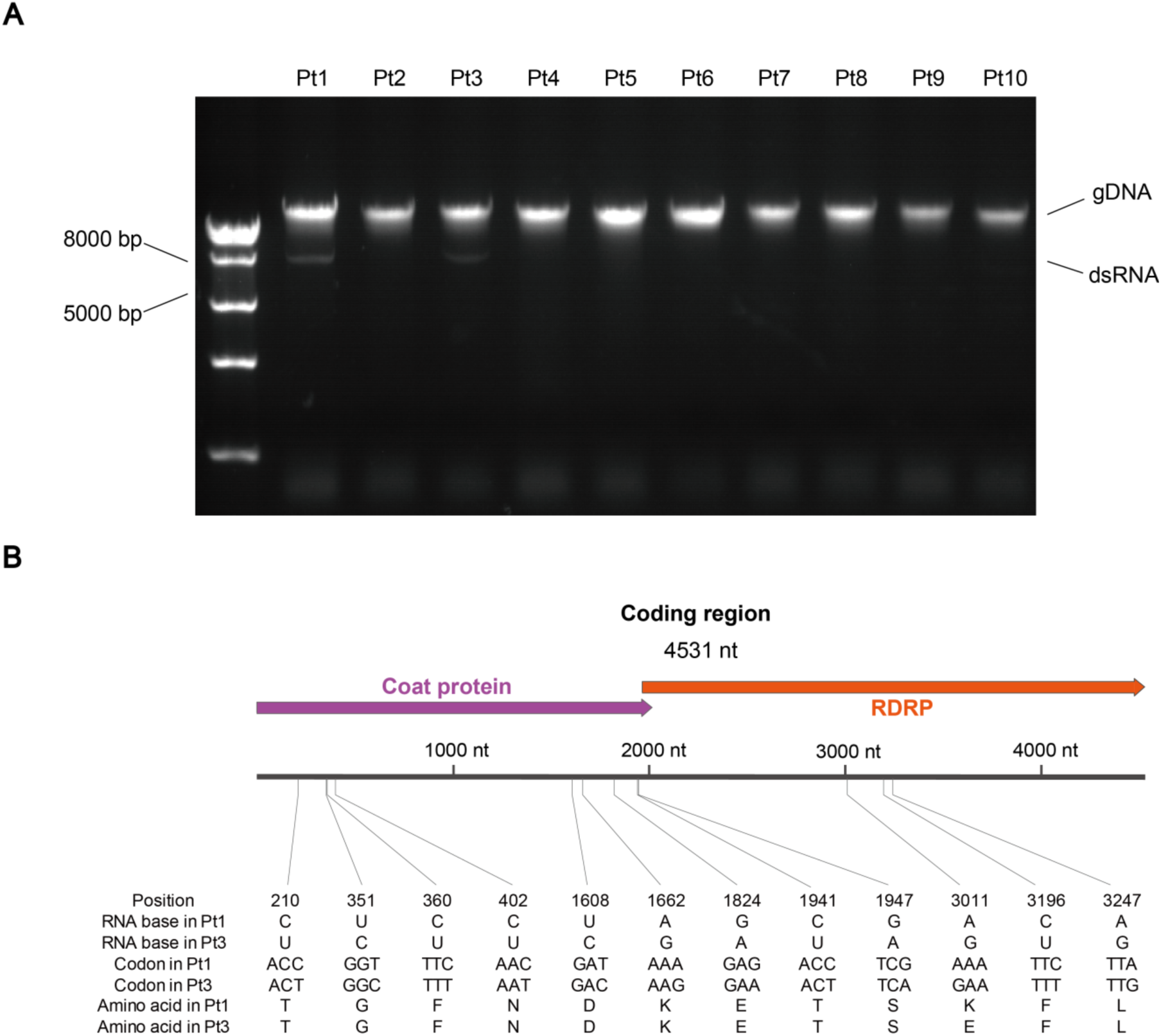
Detection and identification of the dsRNA virus in diatom strains *P. tricornutum* Pt1 (CCMP2561) to Pt10. (A) Agarose gel electrophoresis profile of DNA extracts from Pt1 to Pt10, visualized on a 1% agarose gel. The left numbers indicate the sizes (bp) of the DNA marker, and the right annotations indicate the nucleic acid types for each band or region. (B) Schematic representation highlighting the details of single nucleotide polymorphisms between the coding regions of the dsRNA in Pt1 and Pt3. Note that the Pt1 strain used in this panel is CCMP2561 maintained in Hanhua Hu Lab (see **Materials and Methods**).

To obtain the sequence of the dsRNA-like nucleic acid, we performed strand-specific dsRNA-sequencing as described in the **Materials and Methods** section. We selected the assembled version generated by the Trinity software [35] with the predicted length. We confirmed that both reverse complementary RNAs were detected in the same order (**Fig. 1B**), proving that this nucleic acid is indeed dsRNA. We were unable to find similar sequences in the nuclear or organelle genomes of *P. tricornutum* [16, 45, 46], which argues against the possibility that the dsRNA is generated from *P. tricornutum* DNA transcription.

### Analysis of the detailed sequences of dsRNA Toti-like virus

We then employed RACE (Rapid Amplification of cDNA Ends) to determine the 5’ and 3’ end sequences of both positive and negative strands. PCR products with a consistent length were obtained (**Fig. S1B**). Based on the predominant sequences (**Fig. S1A and Supplementary Table S2**), the negative strand of the virus genome may harbor an additional base at each terminal compared to the positive strand (**Fig. 1C**). Previous studies on marine *Totiviridae* were limited in their ability to determine all the terminal sequences of the dsRNA genome due to constraints in meta-transcriptomic technology [47, 48] or the assumption that the terminal sequences of the positive and negative strands were identical [48, 49]. Our findings highlight the complexity of the genome terminals of this Toti-like virus and suggest the presence of similar complexity in other members of *Totiviridae*.

We performed bioinformatic studies on the viral sequences obtained. Interestingly, highly similar sequences with 99% identities could be found on NCBI database (Sequence ID: OP191686 and OP191687). These two sequences were detected in *Nannochloropsis oceanica* (*N. oceanica*) and *Thalassiosira weissflogii* (*T. weissflogii*) commonly existing in marine environment and predicted to be the genome of a kind of marine Toti-like virus belonging to *Totiviridae* [48]. The positive strand of the dsRNA genome was shown (**Fig. 1D**) and compared with the sequences found in *N. oceanica* and *T. weissflogii* [48]. Our findings strongly suggest that the model diatom strain Pt1 contains a marine Toti-like virus, the hosts of which were diverse and prevalent in marine unicellular microalgae. Marine *Totiviridae* has been reported to be one of the most widespread families in non-vertebrate organisms [47], especially in marine microalgae [48, 49]. Consequently, the discovery of a kind of marine Toti-like virus in model diatom strain Pt1 displayed great importance for deep research on the relationship between the host and marine *Totiviridae*.

We also found that the terminals of each strand of dsRNA in *P. tricornutum* were distinct from the versions in *N. oceanica* and *T. weissflogii* named Taphios ghabri-like virus 1 isolate Thaweiss1 (OP191686) and Taphios ghabri-like virus 1 isolate Nanocea1 (OP191687) [48]. However, the coding regions were almost the same (identity > 99.5%), suggesting that the three hosts possessed different strains of the same virus. These results demonstrate the prevalence of this virus in stramenopile hosts. And these viruses were bioinformatically established as a new *Totiviridae* genus [48].

The conservative region of the dsRNA genome only contained two long overlapped (+1 ribosomal frameshift) open reading frames (ORFs), which was considered as coat protein (CP) and RNA-dependent RNA polymerase (RDRP) (**Fig. 1D**). Such result was corresponding to the previous report in *N. oceanica* and *T. weissflogii* [48]. Accordingly, part of the sequence of the virus was found in a recently published transcriptome (SRR12347810) of Pt1 [50]. It remains undetermined whether this dsRNA virus infected an ancient common ancestor of stramenopiles, or has horizontally spread. The first scenario is supported by the observation that totiviruses are typically vertically transmitted [51] but would require independent events that led to the loss of the virus in many lineages.

### Distribution of Toti-like virus in other *P. tricornutum* strains

To investigate the presence of the Toti-like virus in *P. tricornutum*, we examined the presence of suspected dsRNA in nine additional strains (Pt2 to Pt10) of *P. tricornutum* collected from various global regions [21]. Apart from Pt1, we observed a putative band of the dsRNA in only Pt3 (**Fig. 2A**). Nuclease digestion experiments revealed that the detected nucleic acid was sensitive to RNase A but resistant to DNase I and nuclease S1 (**Fig. S2**), confirming its dsRNA nature. Furthermore, we conducted a comparative analysis of the protein-coding sequence of this dsRNA with that of Pt1, revealing a high level of similarity (identity > 99.5%) (**Fig. 2B**). This finding suggests that Pt1 and Pt3 harbor different variants of this virus. Notably, both Pt1 and Pt3 were collected from the vicinity of the United Kingdom [21]. The virus may have infected their common ancestor or may have spread horizontally.

### Generation and characterization of virus-removed strain isogenic to Pt1

To further advance the establishment of a diatom-virus research model, the depletion and rescue of the virus were crucial steps. However, these steps have not been previously performed in any diatom-virus interaction system. Previous attempts at virus rescue research have often yielded failed cases [52, 53], indicating the undiscovered complexity of *Totiviridae* amplification in hosts. Therefore, further molecular and genetic research on *Totiviridae* was deemed necessary. Until now, research on virus depletion of *Totiviridae* in their hosts has been limited. While the yeast L-A virus, a member of *Totiviridae*, could be depleted using cycloheximide [54], the same strategy proved ineffective for LRV1, another member of *Totiviridae* [39]. Various methods with unstable effects have been reported for LRV1 depletion as well [39, 55, 56]. We sought to generate a virus-removed strain that is isogenic to the virus-bearing Pt1 strain. The nucleotide analogue 2’-C-methyladenosine (2CMA), which was found to be effective in depleting LRV1 in *Leishmania guyanensis* [39], also accomplished virus removal in Pt1 (**Fig. 3B**). However, Remdesivir, which targets the RDRP of certain single-stranded RNA viruses [57, 58], did not show any effects (**Fig. 3B**). After virus depletion, the clones were cultured in liquid f/2 media to detect dsRNA (**Fig. 3A**). Typically, after five rounds of dilution and culture with 2CMA treatment, nearly all clones obtained through sub-cloning on f/2 plates should be depleted of dsRNA (**Fig. 3A**). This result suggests that the amplification of dsRNA in Pt1 relies on its own encoded RDRP and highlights the robustness of 2CMA for virus depletion.

**Fig. 3.**
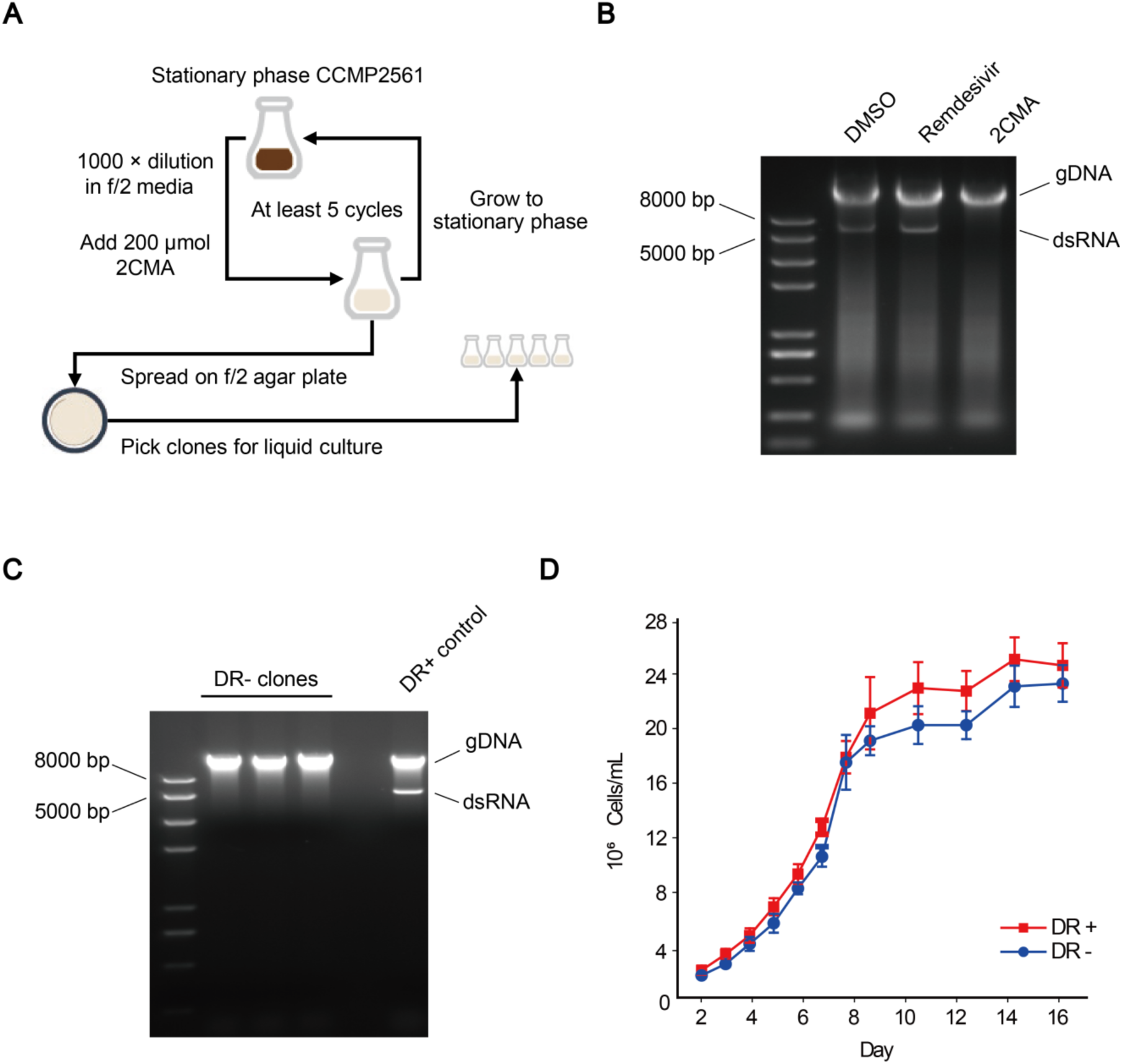
Strategy for generating isogenic, virus-free strains and characterization of virus-containing and virus-free strains. (A) A flow chart illustrating the steps involved in generating isogenic virus-free strains. (B) Agarose gel image displaying DNA samples from Pt1 treated with dimethyl sulfoxide (DMSO), Remdesivir, and 2’-C-Methyladenosine (2CMA). (C) Agarose gel image showing the electrophoretic profile of DNA samples from three DR- clones cultured after 20 rounds of inoculation, compared with the DR+ control. In (B) and (C), the left numbers indicate the sizes (bp) of the DNA marker, and the right annotations indicate the nucleic acid types for each band or region. (D) Growth curves of both DR+ and DR- groups, with 6 biological replicates each grown from a separate colony for each of the two groups.

To ensure the stability of dsRNA depletion, three random clones were selected after the dsRNA depletion process, and they were subjected to 20 rounds of liquid culture (1000-fold dilution in f/2 media) without the addition of 2CMA. All clones maintained the dsRNA-depleted status (**Fig. 3C**), indicating that the dsRNA depletion method (**Fig. 3A**) was reliable and capable of generating relatively stable dsRNA-free clones. By generating isogenic strains of Pt1 with and without this virus (DR+ and DR-), further observations and tests could be conducted. Our results also suggest that 2CMA might be effective in *N. oceanica* and *T. weissflogii* [48], as they are closely related to Pt1, and it was worth exploring its potential in other viruses with closer evolutionary relationships.

The infection by *Totiviridae* can be severe [59] though most hosts of *Totiviridae* are symptomless [51]. We compared the growth curves of six DR+ clones and six DR-clones but found no significant differences (**Fig. 3D**). We tentatively considered this Toti-like virus to be symptomless and temperate under normal laboratory conditions. However, it is difficult to conclude that this virus does not affect the host in the wild. Nonetheless, we speculate that the presence of this virus in the host does not compromise the viability of the host in the marine environment, as microalgae cells infected with obviously toxic viruses do not easily accumulate. Consequently, the viruses found in microalgae clones isolated from the marine environment are mostly temperate. This limited the discovery of virulent marine microalgae viruses significantly hampers in-depth research on them. Avirulent viruses could play important roles in various biological functions of the marine hosts [60]. Regarding this Toti-like virus in *P. tricornutum*, *N. oceanica*, and *T. weissflogii*, it is still challenging to determine whether this virus contributes to the hosts in the wild. However, it is certain that this system represents a mode of interaction between the host and a temperate virus that differs from the previously reported interaction mode [60]. The establishment of such system revealed another mode of temperate infection, contributing to the research on temperate viral infection on marine algae.

### The anti-coat-protein antibody for investigating potential host-virus interactions

The interaction between hosts and *Totiviridae* has been relatively understudied in previous research, primarily due to the typically asymptomatic infections caused by *Totiviridae*. We sought to take advantage of the DR+ Pt1 strain and DR- isogenic strain to probe into this problem. We first generated a rabbit polyclonal antibody against the coat protein for immunoprecipitation studies (**Fig. 4, A and B**). Signals were observed in the DR+ samples, with the most predominant band close to the expected size of the coat protein (73.7 kDa). In contrast, almost no signal was observed for the DR- samples, indicating an excellent specificity (**Fig. 4A**). Using this antibody, we examined the expression of the coat protein during the growth process of Pt1 and observed its relative stability (**Fig. 4C**). This finding suggests that the virus maintains a stable proliferation rate throughout the growth process of *P. tricornutum* in the lab. The successful production of the antibody provides a valuable tool for further molecular-level research on the interaction between the virus and its hosts.

**Fig. 4.**
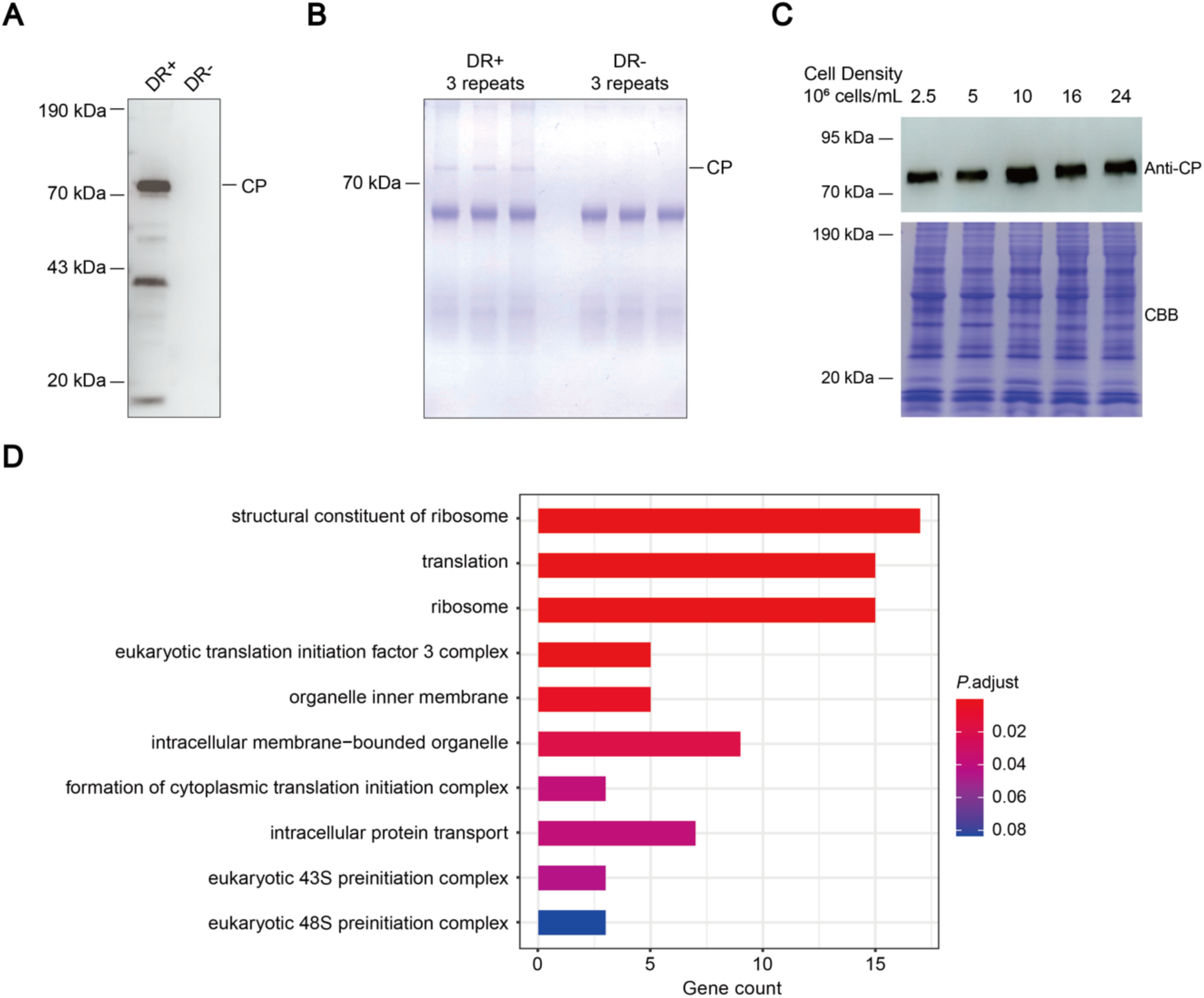
Effect and use of anti-coat-protein antibody. (A) Immunoblot analysis demonstrating the specificity and effectiveness of the anti-coat-protein antibody. (B) SDS-PAGE gel image stained with Coomassie Brilliant Blue G-250, showing the immunoprecipitation sample using the anti-coat-protein antibody from DR+ and DR- total protein. In (A) and (B), CP refers to coat protein. (C) Immunoblot analysis depicting the expression of coat protein during the growth process of Pt1, with the loading control provided by the SDS-PAGE gel image stained with Coomassie Brilliant Blue G-250. In (A), (B), and (C), the left numbers indicate the sizes (bp) of the protein marker, and the right annotations indicate the bands of full-length coat protein. In (C), the numbers on the top indicate the cell density when collecting the samples. (D) Gene ontology (GO) analysis of the candidate list of host proteins interacting with the virus coat protein, obtained from mass spectrometry data of co-immunoprecipitation using the anti-coat-protein antibody. The bar chart displays the top 10 representative GO terms, with the X-axis representing the number of genes in each GO term.

To further elucidate the potential viral influence on the host at the molecular level, we performed immunoprecipitation using the anti-coat-protein antibody followed by mass spectrometry. This approach allowed us to generate a list of proteins that may interact with the coat protein (**Supplementary Table S4**). Among the candidates, 187 proteins are nuclear-encoded and 148 of them are annotated with gene ontology (GO) terms. Notably, we observed an enrichment of translation-related proteins associated with GO terms such as “structural constituent of ribosome” (GO:0003735), “translation” (GO:0006412), “ribosome” (GO:0005840), “eukaryotic translation initiation factor 3 complex” (GO:0005852), and “eukaryotic 43S preinitiation complex” (GO:0016282) (**Fig. 4D and Supplementary Table S5**). This suggests that the coat protein likely interacts with ribosomes and translation initiation-related proteins. We also observed an enrichment of membrane-related proteins, represented by GO terms such as “organelle inner membrane” (GO:0019866), “intracellular membrane-bounded organelle” (GO:0043231), and “intracellular protein transport” (GO:0006886) (**Fig. 4D and Supplementary Table S5**). We hypothesize that some coat proteins may interact with ribosome-related proteins in close proximity to various membrane structures (**Fig. 4D and Supplementary Table S6**).

Based on the results of co-immunoprecipitation, we speculate that the presence of this virus in *P. tricornutum* may cause instability or remodeling in host translational processes, intracellular protein transport, and some unclear certain membrane-dependent processes. Our findings highlight the potential risks associated with molecular-level research using the model strain Pt1.

## Discussion

Our study successfully identified a dsRNA virus belonging to the *Totiviridae* family in the Pt1 diatom strain, which is one of the early-sequenced diatom strains. Additionally, we discovered a geographically similar sampling location where the Pt3 strain of *P. tricornutum* also harbors this virus. It is worth emphasizing the risks associated with relying solely on the Pt1 or Pt3 strain for studying specific processes and underscores the importance of utilizing multiple accessions of *P. tricornutum* [20, 21, 23].

In addition, our study provides a valuable strategy for obtaining virus-free *P. tricornutum* clones, which allows for the exclusion of the virus’s effects on Pt1 and facilitates more rigorous investigation in these risky research fields. By employing a system comprising both virus-containing and virus-free strains, we confirmed the latent infection of this virus. Additionally, we successfully generated a highly effective anti-coat-protein antibody, serving as a valuable tool for molecular research. Our investigations revealed several potential aspects affected by this virus on the host, including translation and intracellular membrane-related mechanisms.

Furthermore, this study highlights a novel case of latent infection in diatom hosts, which has not been previously reported. Future research on this virus can focus on detailed understanding of its interactions with host cells, such as its maintenance requirements and the possibility of infecting virus-free cells using extracted viral RNA. Such studies may lead to the emergence of novel transgenic tools for diatoms.

## Author contributions

Jiahuan ZHANG, Hanhua HU, and Xiaobo LI conceived and designed the study. Jiahuan ZHANG, Chenjie LI, Xiaofeng XIONG, and Huan ZHANG conducted the physiological, biochemical, and molecular experiments. Kangning GUO, Yanlei FENG, and Jiahuan ZHANG conducted bioinformatic analysis. Jiahuan ZHANG and Xiaobo LI wrote the manuscript with input from all authors.

## Acknowledgements

We thank Chris Bowler at École Normale Supérieure for providing *P. tricornutum* strains. In addition, we thank Jin HU, Jia CHEN, Xue BAI, Mingzhu FAN and Shan FENG in the Mass Spectrometry & Metabolomics Core Facility (MSMCF) at Westlake University for assistance in mass spectrometry experiments. This research is supported by a Zhejiang Province Key R&D Program grant (2023SDXHDX0002), a National Key R&D Program of China grant (2019YFA0906300), National Natural Science Foundation of China grants (42106114, 32170255), and a Westlake Education Foundation grant (WU2023B002).

## Supplementary Figures

**Fig. S1.**
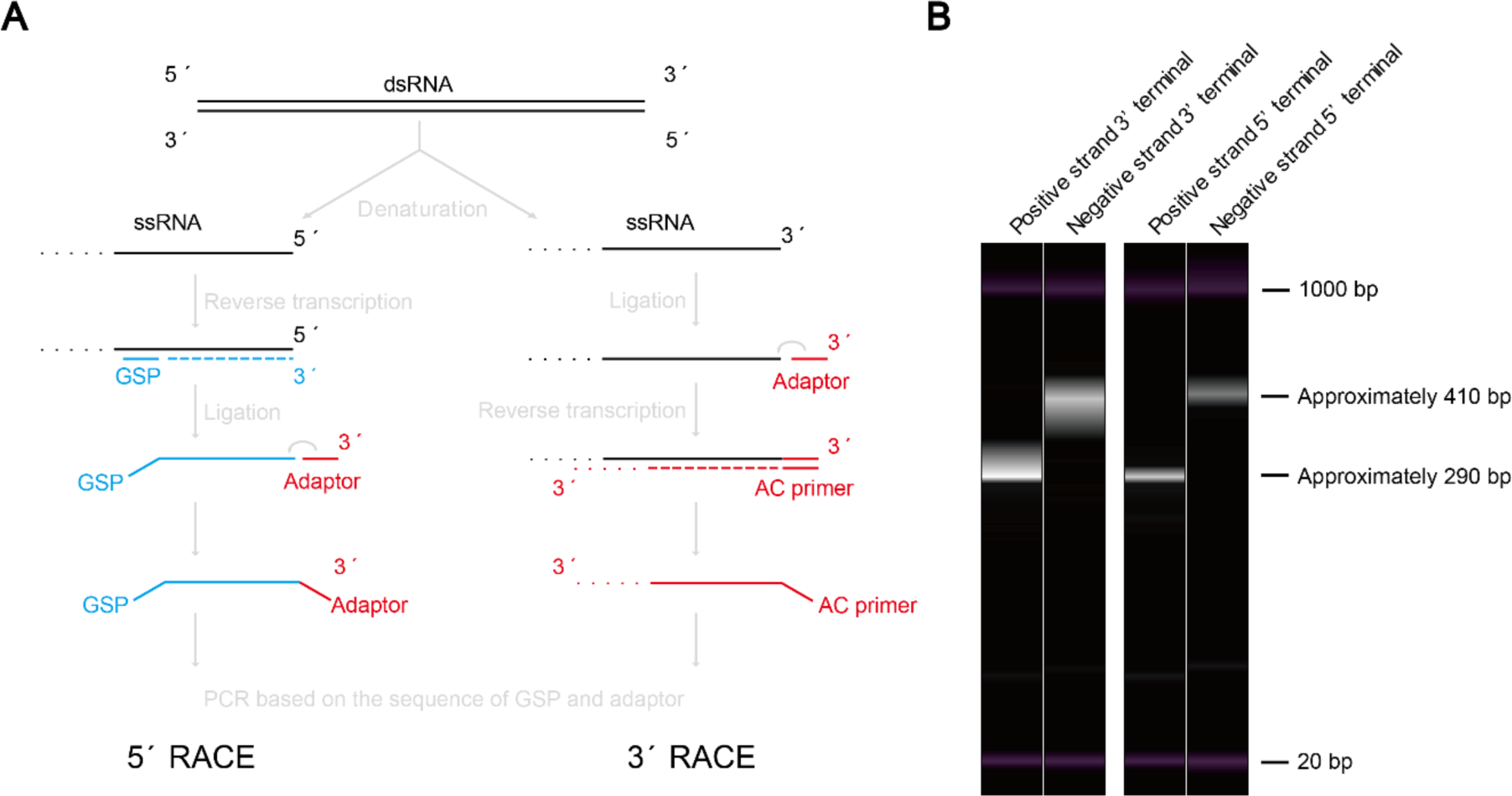
The strategy and the results of RACE (Rapid Amplification of cDNA Ends) library construction for next-generation sequencing. (A) A flow chart illustrating the steps involved in RACE experiment for both 5’ and 3’ terminal sequences of the viral dsRNA genome. (B) Simulated gel images displaying the libraries of RACE for next-generation sequencing, analyzed on Qsep100. See **Materials and Methods** for detailed steps of RACE and **Supplementary Table S2** for detailed results of RACE.

**Fig. S2.**
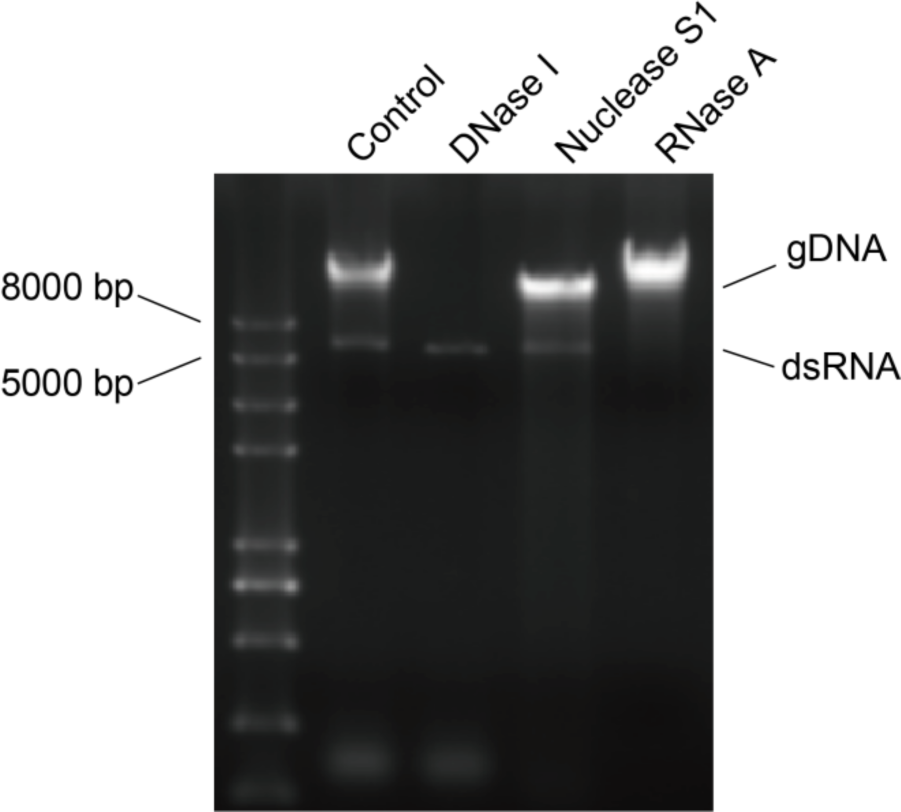
Agarose gel image displaying the electrophoretic profile of DNA extracts from Pt3 treated with DNase I, Nuclease S1, or RNase A, resolved on a 1% agarose gel.

